# Varied monoamine reuptake inhibitors reduce parvalbumin expression; implications for pyramidal cell disinhibition and enhanced neuroplasticity

**DOI:** 10.64898/2025.12.12.693967

**Authors:** Katie Hummel, Giorgi Shautidze, Eric Thorland, Mathew Amontree, Pauline Wonnenberg, Ismary Blanco, Katherine Conant

## Abstract

First-line antidepressants are effective in a significant percent of individuals but a full understanding of how these therapeutics target specific endpoints is lacking. Prior work has shown that depression is associated with hippocampal atrophy and that antidepressants can increase neurotrophin levels to increase hippocampal neurogenesis as well as hippocampal pyramidal cell (PC) spine density and arbor. These effects likely contribute to amelioration of symptoms. A less well-explored possibility is that antidepressants concomitantly disinhibit hippocampal PC activity, which could also facilitate increased PC arbor, spinogenesis and/or activity. In accordance, previous studies have shown antidepressants can attenuate stress-induced upregulation of perineuronal nets (PNNs). PNNs are predominantly localized to parvalbumin (PV) expressing GABAergic interneurons and increase PV expression and neuronal activity. Though specific antidepressants have been explored for effects on regional PNN expression, the question of whether hippocampal PNN/ECM remodeling is a shared feature of varied antidepressant drugs and more importantly, of whether it is associated with significant hippocampal PV inhibition, has not been well-addressed. Herein we examine three monoamine reuptake inhibitors, fluoxetine, venlafaxine and viloxazine, in animal models for effects on PNN remodeling and PV expression, a proxy for PV activity. We observe shared effects of these therapeutics including the ability to increase PNN degrading effectors that can downregulate PV activity. Consistent with this, we observe shared effects of these drugs in terms of their ability to significantly reduce PV levels. These findings highlight the possibility that ECM remodeling and associated hippocampal PC disinhibition represent a shared feature of varied antidepressant medications.

## Introduction

Major depressive disorder (MDD) has been linked to deficits in overall cortical and hippocampal excitatory neuronal activity, and effective antidepressant treatments including monoamine reuptake inhibitors and electroconvulsive therapy (ECT) generally increase excitatory activity. At the structural level, ECT and monoaminergic modulators have been shown to increase hippocampal neurogenesis, spinogenesis and pyramidal cell (PC) arborization [1-3], and at the molecular level monoaminergic drugs can increase the phosphorylation and membrane insertion of glutamate receptor subunits [4, 5]. A non-mutually exclusive mechanism that could enhance PC excitability includes reduced PV activity and thus diminished PV-mediated PC inhibition. This is supported by the ability of ketamine, which is somewhat selective in its ability to inhibit NMDA receptors expressed on PV neurons, to rapidly improve mood.

Perineuronal nets (PNNs) are a dense form of extracellular matrix that predominantly surrounds PV-expressing neurons in cortical areas and dorsal hippocampus. During development, the maturation of PNNs is associated with closure of critical periods of neuronal plasticity [6-8]. Recent work, however, has shown that PNN deposition may be further increased in adulthood by aging, inflammation and stress [9], including chronic stress from paradigms linked to depression-relevant behavior in rodents [10-12]. In addition, PNNs can also be attenuated in adulthood in association with gamma entrainment, ketamine and other antidepressant medications [12-14].

PNN levels in turn can have a significant influence on PV activity. Recent work has shown that PNNs can increase PV cell membrane capacitance, restrain lateral diffusion of glutamate receptors and potentially enhance GluA receptor subunit insertion [15-17]. These effects could in turn contribute to increased PV activity, and thus PC inhibition, in conditions associated with reduced PNN cleavage and/or increased PNN deposition [10-12]. Of relevance to MDD, our lab has previously observed that in a chronic corticosterone murine model of depression PNN intensity is increased [12], and another group has shown that PNN component levels are also increased in rats following exposure to chronic stress [11].

In previous work we observed that venlafaxine, a serotonin norepinephrine reuptake inhibitor, could attenuate PNNs in an MMP-9 dependent manner to increase gamma oscillation power [12, 18], which is reduced in major depression [19]. While gamma power may be increased in association with increased excitatory/inhibitory balance and/or PC disinhibition [20, 21], prior studies have not explored more cell-type specific correlates of widespread PC disinhibition with venlafaxine. In addition, to address the question of whether PC disinhibition may be a shared mechanism of varied antidepressants, we compared effects of venlafaxine to other monoamine reuptake inhibitors for the ability to concomitantly stimulate PNN remodeling and PC disinhibition in mice or zebrafish.

## Materials and Methods

### Animal subjects

Wild-type late juvenile/early adult (2-4 months) zebrafish with a 50:50 female:male ratio were housed in groups of 10-20 in 2L tanks with water temperature kept at 28°C. Zebrafish were fed either once or twice a day and were maintained on a 14/10 light-dark cycle (lights on 9 AM and lights off at 11 PM). All procedures were performed in accordance with the Institutional Animal Care and Use Committee of Georgetown University, Washington DC, USA (Protocol # 2020-0034). Euthanasia for zebrafish was performed following tricaine methane sulfonate anaesthesia and decapitation.

6 week old female C57BL/6 mice were purchased from Jackson laboratories and housed 3 mice per cage with enrichment dens or balconies. All procedures were performed in accordance with the Institutional Animal Care and Use Committee of Georgetown University, Washington DC, USA (Protocol # 2018-0037). Euthanasia for mice was performed with deep isoflurane anaesthesia (inhalation) followed by confirmation of a lack of response to deep pain and rapid decapitation.

### Monoamine reuptake inhibitors

Fluoxetine (Sigma; CAS# 56296-78-7) treatment of zebrafish was based on the dosage used in a prior publication [22] or 100μg/L, and to control for drug degradation, fish in control and treatment groups were netted and moved to new tanks with fresh vehicle or fluoxetine solution every two days during treatment. Separate cohorts were used for proteomics and novel tank behavior, but for both cohorts the treatment duration was for 35-37 days.

Venlafaxine (30 mg/kg/day) and viloxazine (50 mg/kg/day) treatment of mice was administered with once daily intraperitoneal (IP) injections, on alternate sides each day, in 200 μl saline. Venlafaxine and viloxazine were also ordered from Sigma and dissolved in saline and then sterile filtered into aliquots. Control mice were injected once daily with sterile saline. Mice began treatment at approximately 6 weeks and treatment duration was 40 days. The elevated plus maze experiment was performed on treatment day 15.

### Proteomics and ELISA

Proteomic analyses were performed as previously described for zebrafish and murine samples, using our core facility run by Dr. Junfeng Ma [23-25]. 100 µg of total protein from individual murine brain or fish telencephalon lysates was sent for proteomics analysis using the Orbitrap Lumos Tribrid mass spectrometer instrument.

Proteins in pellets were extracted by using lysis buffer containing 10% SDS, 1x proteinase inhibitor cocktail and 50 mM triethylammonium bicarbonate. After the addition of benzonase, the cellular suspension was incubated on ice for 20 min followed by sonication with a probe-tip sonicator for 5 pulse (10 sec on 20 sec off for each pulse). The cell lysate was centrifuged for 15 min, 16000 g at 4 °C, with the supernatant saved and used for protein concentration determination by BCA assay. Equal amount of proteins from each sample was then treated with DTT and iodoacetamide and then loaded onto a S-Trap column (ProtiFi, LLC) by following the manufacturer’s instructions. Proteins were digested with sequencing-grade Lys-C/trypsin (Promega) by incubation at 37°C overnight. The resulting peptides were eluted and dried down with a SpeedVac (Fisher Scientific). To construct a comprehensive spectral library, an aliquot of each digest was pooled and fractionated by high pH reversed phase fractionation into 8 fractions, peptides in each fraction were dried down with a SpeedVac.

Peptides were analyzed with a nanoAcquity UPLC system (Waters) coupled with Orbitrap Fusion Lumos mass spectrometer (Thermo Fisher). Samples in 0.1% FA solution are loaded onto a C18 Trap column (Waters Acquity UPLC M-Class Trap, Symmetry C18, 100 Å, 5 μm, 180 μm x 20 mm) at 10 µL/min for 4 min. Peptides are then separated with an analytical column (Waters Acquity UPLC M-Class, peptide BEH C18 column, 300 Å, 1.7 μm, 75 μm x 150 mm) with the temperature controlled at 40°C. The flow rate is set as 400 nL/min. A 150-min gradient of buffer A (2% ACN, 0.1% formic acid) and buffer B (0.1% formic acid in ACN) is used for separation: 1% buffer B at 0 min, 5% buffer B at 1 min, 22% buffer B at 90 min, 50% buffer B at 100min, 98% buffer B at 120 min, 98% buffer B at 130 min, 1% buffer B at 130.1 min, and 1% buffer B at 150 min. Data files were acquired with data independent acquisition (DIA) mode or data dependent acquisition (DDA) on an Orbitap Fusion Lumos mass spectrometer using an ion spray voltage of 2.4 kV and an ion transfer temperature of 275°C. Mass spectra were recorded with Xcalibur 4.0.

Analysis of DDA raw files was performed in Proteome Discoverer (Thermo Fisher Scientific, version 2.4) with Sequest HT database search engines. The database for Mice or Zebrafish was downloaded from Uniprot. The database-searching parameters were set as below: full tryptic digestion and allowed up to two missed cleavages, the precursor mass tolerance was set at 10 ppm, whereas the fragment-mass tolerance was set at 0.02 Da. Carbamidomethylation of cysteines (+57.0215 Da) was set as a fixed modification, and variable modifications of methionine oxidation (+15.9949 Da), acetyl (N-terminus, +42.011 Da) were allowed. The false-discovery rate (FDR) was determined by using a target-decoy search strategy. The decoy-sequence database contains each sequence in reverse orientations, enabling FDR estimation. On the peptide level, the corresponding FDR was less than 1%. Analysis of DIA raw files was done by using Spectronaut (Biognosys, v15) with hybrid library built from 8 DDA data files (from high pH reversed phase fractionation) and 10 DIA data files.

ELISA was performed using an IL-33 ELISA kit from R and D (Minneapolis MN) which was performed according to the manufacturer’s instructions.

### Behavioral assays

#### Novel Tank Test (Fish)

Long cylindrical tanks were filled to 8 liters with normal tank water. Fish were placed in the tank by inverting net and lightly tapping on the edge of the tank. Recording time started when the fish entered the water. The tanks were dimly lit with red light to reduce glare and add novelty to the environment. Fish were recorded for an hour. AnyMaze cameral tracking software was used to analyze which “level” of the tank the fish spent time in. Each level was drawn as a multiple of n, increasing with depth. That is level one had a depth of n (Top), level two was a depth of 2n (Middle Top), level three was a depth of 3n (Middle Bottom) and so forth.

#### Elevated Plus Maze (Mice)

The elevated plus maze was performed using AnyMaze video tracking software (black mice on a white apparatus were tracked), using a previously described protocol [12].

### Statistics

Statistical tests were done using GraphPad Prism. Data outlier analyses were performed using the Grubs outlier test. Unpaired Student’s *t* tests were used to compare 2 group data where indicated. Statistics for volcano plot data were performed as described [25].

## Results

### I. Varied monoamine antidepressants modulate brain lysate levels of PNN effectors including IL-33 and cathepsins

In previous work, with targeted Western blot and ELISA analyses, venlafaxine increased cortical and hippocampal MMP-9 levels in both non-treated and chronic corticosterone treated C57/bl6 mice [12]. Moreover, humans with MDD who were on monoamine modulating medications including selective serotonin and serotonin/norepinephrine reuptake inhibitors showed an increase in prefrontal cortex MMP-9/tissue inhibitor of MMPs-1 (TIMP-1) ratio [12]. In the current study we have utilized proteomic analyses to identify additional targets of the serotonin norepinephrine reuptake inhibitor (SNRI) venlafaxine and the selective norepinephrine reuptake inhibitor viloxazine in C57/bl6 mice. We also studied effects of the selective serotonin reuptake inhibitor (SSRI) fluoxetine using a zebrafish model.

Results are shown in figure 1. As can be appreciated, proteomic analyses of zebrafish telencephalon lysates showed changes in ECM effectors following treatment with fluoxetine. In particular, levels of cathepsin La and cathepsin F were elevated. Though they have intracellular substrates, cathepsins can be secreted into the extracellular space where they remodel the ECM and also activate additional ECM targeting proteases including MMPs [26, 27]. Proteomics analyses in mice demonstrate that venlafaxine increased levels of A disintegrin and metalloproteinase with thrombospondin repeats-10 (ADAMTS-10), which can reduce profibrotic TGF-β signaling in brain and retina, while venlafaxine reduced levels of serine protease inhibitor A3M, which inhibits cathepsin G [28]. Recent work also suggest that select ADAMTS family members can cleave the PNN component brevican [29]. Importantly, both venlafaxine and viloxaxine increased levels of the cytokine IL-33 in murine hippocampus as indicated by ELISA-associated differences for the former and proteomics differences for the latter (Figure 1). Importantly, IL-33 instructs hippocampal microglial engulfment of the PNN component aggrecan to promote dendritic spine plasticity [30].

**Figure 1.**
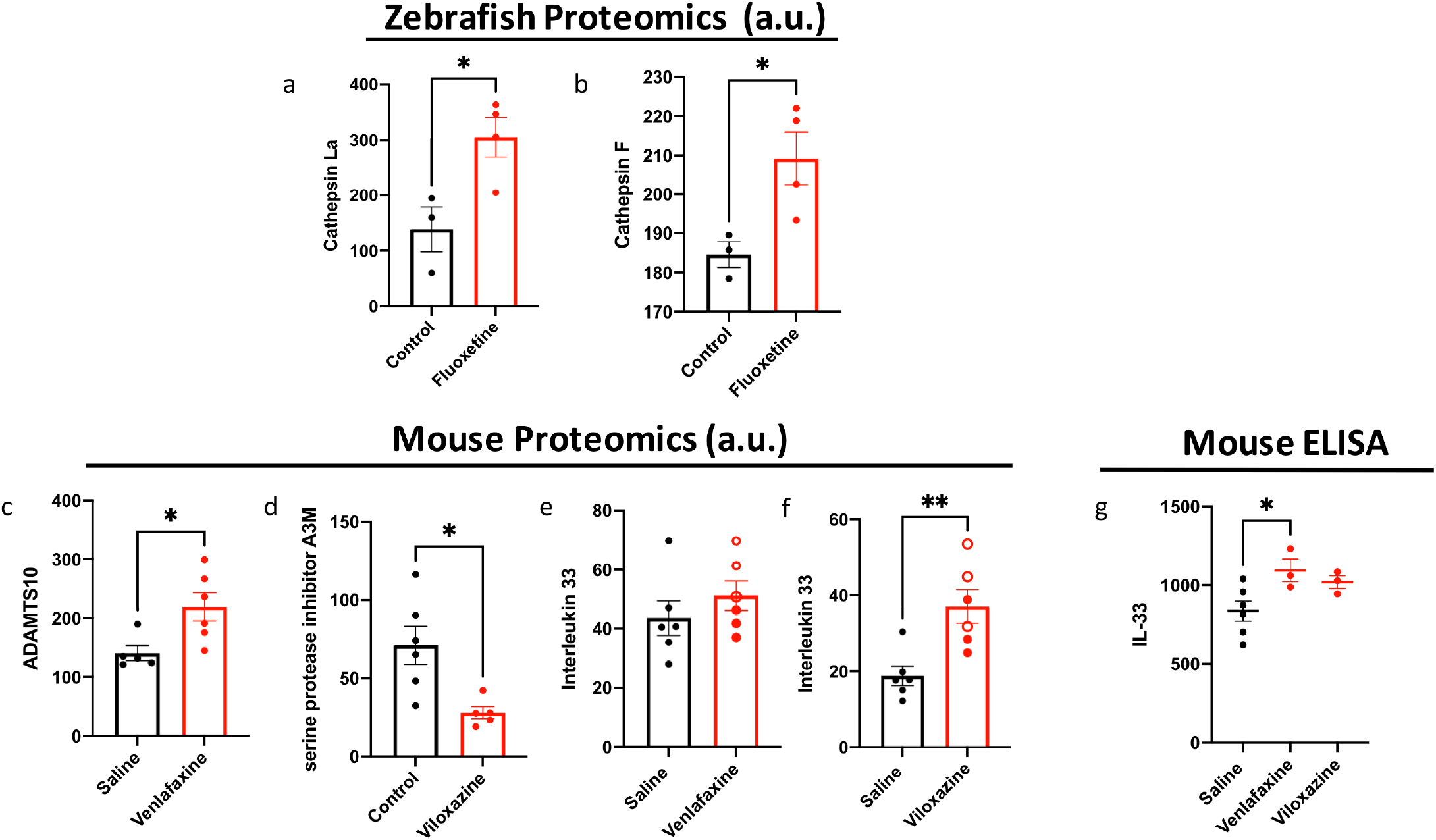
Proteomic and ELISA analyses of zebrafish telencephalon and murine hippocampal lysates show changes in ECM effectors following treatment with varied monoamine reuptake inhibitors. Proteomic analyses of telecephalon lysates from zebrafish show increased levels of cathepsin La (1a; *p*= 0.027) and cathepsin F (1b; *p*= 0.033), as determined by Student’s *t*-test of proteomics values, after treatment with fluoxetine. Analyses of raw data from murine hippocampi demonstrate that venlafaxine increased levels of <u>A d</u>isintegrin <u>a</u>nd <u>m</u>etalloproteinase with thrombo<u>s</u>pondin repeats-10 (ADAMTS-10) (1c; *p*= 0.023, Student’s *t*-test), which can reduce profibrotic TGF-β signaling in brain and retina, and that venlafaxine also reduced levels of serine protease inhibitor A3M (1d; *p*= 0.013), which inhibits cathepsin G [28]. Importantly, venlafaxine and viloxaxine both increased levels of the cytokine IL-33 in murine hippocampus as indicated by proteomics-associated differences for viloxazine (1f; *p*= 0.005) and ELISA-associated differences for the latter (1g; *p*=0.042). Importantly, IL-33 instructs hippocampal microglial engulfment of the PNN component aggrecan to promote dendritic spine plasticity [30]. Note that for ELISA analyses, the proteomics samples indicated by an unfilled circle had been depleted and thus were not included in the ELISA assay.

### II. Monoamine antidepressants reduce levels of PNN components including aggrecan and versican

Aggrecan is a PNN component that is highly expressed in the hippocampus and often used in combination or in lieu of WFA [31]. There is also an increased abundance of WFA-negative, aggrecan-positive PNNs in hippocampus [32], and aggrecan is more likely to be localized to PV neurons than are other PNN components such as brevican [33]. Most important, however, is published work showing that even partial aggrecan attenuation in murine visual cortex leads to reduced parvalbumin expression as well as reinstatement of critical period like plasticity [34]. Indeed there are bidirectional relationships between aggregan expression and PV expression [35, 36].

As shown in figure 2, proteomics analyses showed significantly reduced aggrecan levels in murine hippocampal and zebrafish telencephalon lysates following varied antidepressant drug treatment. Zebrafish lysates also showed reduced levels of versican, an additional PNN component.

**Figure 2.**
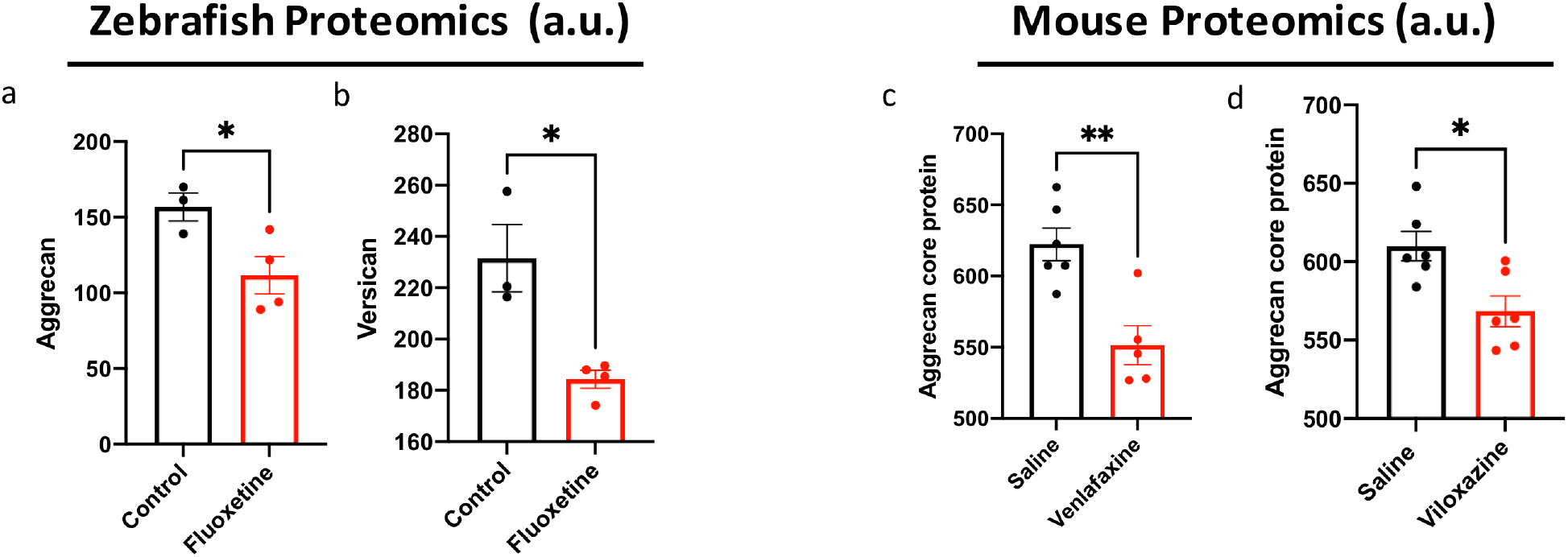
Varied monoamine reuptake inhibitors reduce levels of PNN components including aggrecan and versican. As shown in figure 2, proteomics analyses showed significantly reduced aggrecan levels in zebrafish telencephalon (2a; *p*= 0.042, Student’s *t*-test) and murine hippocampal lysates (2c; *p*= 0.003 and 2d; *p*= 0.012, Student’s *t*-test) following varied monoamine reuptake inhibitor treatment. Zebrafish lysates also showed reduced levels of versican (2b; *p*= 0.010, Student’s *t*-test), an additional PNN component.

### III. Monoamine antidepressants reduce hippocampal PV levels, a proxy for PV activity

Attenuation of PNNs in visual cortex by chondroitinase injection or in hippocampus following chronic venlafaxine administration has been associated with increased gamma oscillation power and/or restoration of juvenile plasticity [21, 37]. Based on electrophysiological recordings as well as published studies examining effects of PNNs on PV neuron membrane capacitance and glutamate receptor insertion/localization, studies of PNN levels and gamma power suggest that PNN disruption reduced PV mediated inhibition. To explore whether varied antidepressants could reduce PV mediated inhibition in the hippocampus, we focused on the ability of these drugs to reduce PV levels. Prior studies have indeed suggested that PV levels and intensity can serve as proxies for the excitability and activity of PV neurons [38, 39]. PV expression is upregulated with increased activity of PV neurons and PV levels are generally increased with increased PNN levels in a cross species manner [39]. As shown in figure 3, proteomic analyses demonstrated that PV levels were significantly reduced in mice and in zebrafish following their treatment with diverse monoamine reuptake inhibitors.

**Figure 3.**
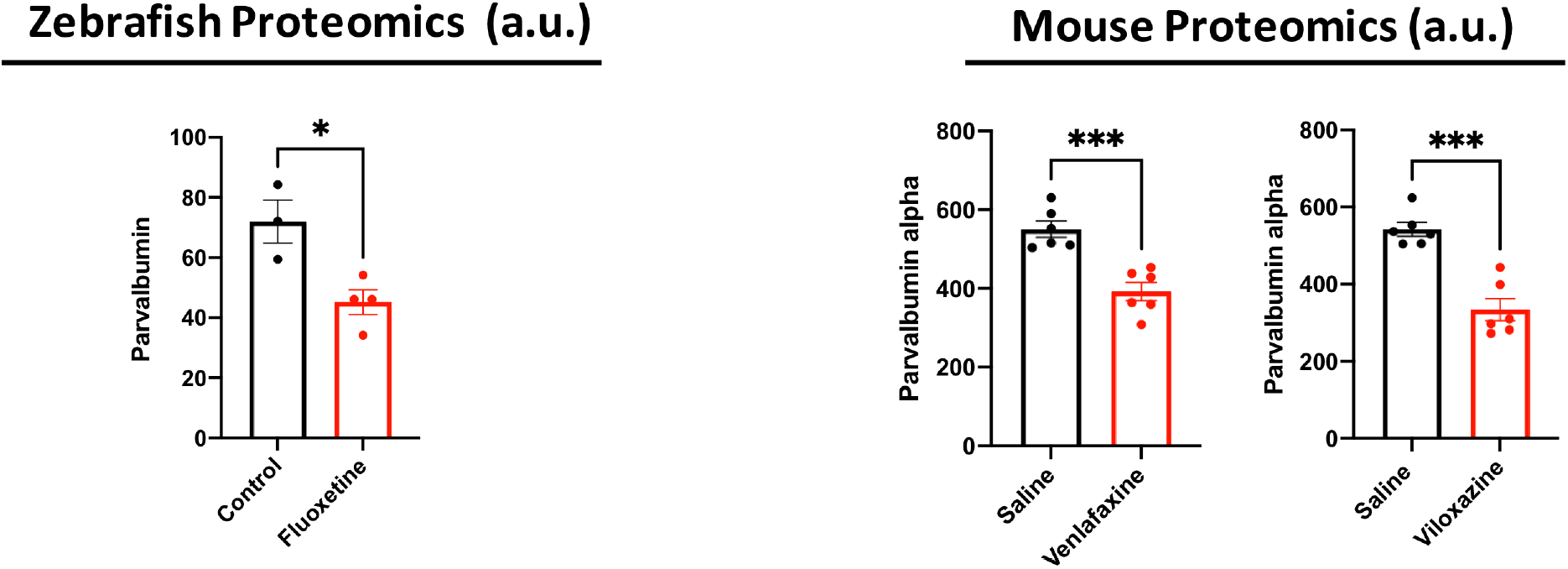
Varied monoamine reuptake inhibitors reduce hippocampal PV levels. To explore whether varied antidepressants could reduce PV mediated inhibition in the hippocampus, we focused on the ability of these drugs to reduce PV levels. Prior studies have indeed suggested that PV levels and intensity can serve as proxies for the excitability and activity of PV neurons [38, 39]. As shown in figure 3, fluoxetine reduced parvalbumin levels in zebrafish telencephalon (3a; *p*= 0.018, Student’s *t*-test) and venlafaxine and viloxazine reduced parvalbumin levels in murine hippocampi (3b; *p*= 0.0005 and 3c; *p*= 0.0001, Student’s *t*-test).

### IV. Proteomic analyses

Though mass spectrometry proteomics analyses can underestimate the magnitude of protein changes due to factors including ratio compression and dynamic range limitations [40], these analyses represent a powerful tool to examine samples in a relatively unbiased manner. Shown in figure 4a is a volcano plot for fluoxetine (zebrafish) as well as venlafaxine and viloxatine treated (mice). Parvalbumin is decreased with each of the three monoamine reuptake inhibitors, and PNN effectors and components of interest, including cathepsin La (ctsla), IL-33, Serpina3m, and aggrecan, are highlighted for select treatments as indicated (fluoxetine in lavender, venlafaxine in orange and viloxazine in pink). Changes that were significant in the Spectronaut analyses, with an -log10(adjusted P-value) of 1.3 and a +/-0.5 log2 fold change, included parvalbumin for each of the 3 drugs, as well as cstla and acanb for fluoxetine and IL33 and serpin3m for viloxazine. In figure 4b heat maps for the top 25 differentially regulated proteins are shown for each group.

**Figure 4.**
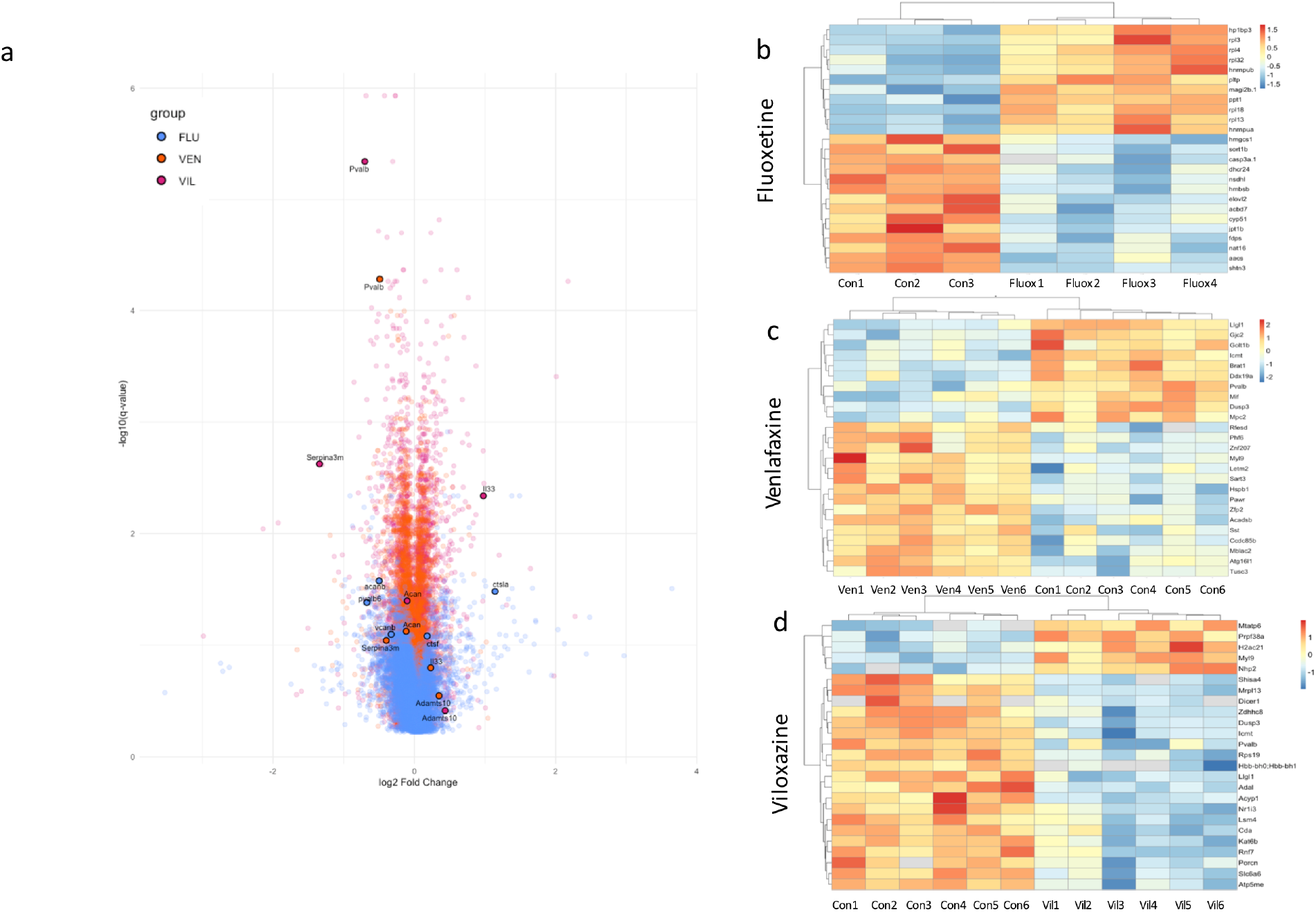
Proteomic analyses show reduced parvalbumin and altered ECM proteins and/or effectors in association with varied monoamine reuptake inhibitors. Shown in figure 4a is a volcano plot for fluoxetine (zebrafish) as well as venlafaxine and viloxatine treated (mice). Parvalbumin is decreased with each of the three monoamine reuptake inhibitors, and PNN effectors and components of interest, including cathepsin La (ctsla), IL-33, Serpina3m, and aggrecan, are highlighted for select treatments as indicated (fluoxetine in lavender, venlafaxine in orange and viloxazine in pink). In figure 4b heat maps for the top 25 differentially regulated proteins are shown for each group.

## Discussion

As described in the present manuscript, we observe ADD associated changes in PNN targeting proteases and PNN components in hippocampal or telencephalon lysates prepared from murine or zebrafish brain. Initial changes were detected by unbiased protein mass spectrometry and further validated with ELISA assays depending on sample availability. Though fish and mouse studies started as separate rotation projects, we were intrigued by the ability of the three monoamine reuptake inhibitors, with differential effects on reuptake inhibition of serotonin or norepinephrine, increase ECM targeting proteases, reduce aggrecan levels, and reduce parvalbumin expression in a cross species manner.

In a previous study, we demonstrated that venlafaxine could increase MMP-9 levels in murine hippocampus and cortex [12]. In addition, venlafaxine treatment was associated with an MMP-9 dependent increase in brevican cleavage [12]. MMP-9 is released from neurons to enhance neuroplasticity through multiple mechanisms [8, 18, 41, 42], and it is also released from microglia to potentially remodel PNNs and synapses. Monoamine that target Gα_s_ coupled GPCRs in particular, such as the β-adrenergic receptor, dopamine 1 receptor and 5-HT7 receptor, have in fact been linked to increased MMP-9 activity and/or release from neurons [12, 43, 44]. We have also reported that MMP-9 levels were decreased in the more anxious of two mouse strains, with the more anxious strain having been reported to have lower levels of serotonin [45, 46]. Importantly, MMP-9 can degrade ECM components, including aggrecan [47], that can regulate PV expression. Of relevance, a recent study using Translating Ribosome Affinity Purification showed that fluoxetine treatment of mice could reduce PNN staining and PV expression in select regions of the murine prefrontal cortex [48].

Data described herein using unbiased proteomics suggests that varied antidepressants may upregulate several ECM effectors mice and fish. Monoamine reuptake inhibitor associated changes of interest detected by proteomics included cathepsins in fish and IL33 in mice. As previously stated in the results section, IL33 instructs microglial engulfment of aggrecan [30]. A caveat of our study is that the lysates we prepared for proteomics did not include isolation of triton X-100 insoluble proteins, which include ECM-bound MMPs [49], and thus we could have missed detecting MMP family members that can also reduce PNN levels in varied contexts [50]. Future studies should include attention to this detail.

Another caveat of our study is that mice and fish were not exposed to a stress paradigm known to increase anxiety and depressive like behaviors. However, we note that our animals were likely stressed in that mice had daily injections and fish had to be regularly netted and transferred to new tanks with freshly added fluoxetine. In addition, previous studies identifying biochemical mechanisms and depression related behavioral effects of fluoxetine and other monoamine modulating antidepressants did not include additional stressors [37, 51].

Data described herein also show a shared ability of the antidepressants to reduce aggrecan levels in mice and fish. Aggrecan is an important component of PNNs in varied species and abundantly expressed in hippocampus. Moreover, a single cell transcriptomics study suggests that PV interneurons are a principal source of the protein [33]. Previously published work has shown that even partial aggrecan attenuation in murine visual cortex leads to reduced parvalbumin expression as well as reinstatement of critical period like plasticity [34]. Bidirectional relationships between aggregan expression and PV expression have also been described [35, 36]. Of interest is a recent study of chronic fluoxetine (21 days) in male mice that reduced aggrecan-positive PNNs in select hippocampal and somatosensory regions [52].

In the present study we also observe reduced PV levels with the three separate monoamine reuptake inhibitors, which is a proxy for reduced PV activity [39]. This was also intriguing in terms of its manifestation in a cross-species and diverse monoamine-reuptake inhibitor dependent manner, as well as its potential importance to E/I balance. Each PV neuron has the potential to inhibit many pyramidal cells, and reduced pyramidal excitation is thought to underlie select deficits observed with major depression [4]. Consistent with a role for disinhibition in antidepressant efficacy, the rapidly acting antidepressant ketamine can also disinhibit pyramidal neurons in that it preferentially inhibits GluN receptors on PV neurons [53, 54]. In addition, sub-chronic treatment with ketamine also reduces WFA labeled PNNs [13]. Moreover, sub-anesthetic doses of ketamine increase gamma oscillation power in a manner consistent with the disinhibition hypothesis [55]. Importantly, gamma power is reduced in human with MDD and animal models of the same [19, 56, 57]. Of interest, however, is that selective activation of 5HT2A subtype serotonin receptors in ventral hippocampus may instead stimulate acute activation of PV interneurons and reduced anxiety-relevant behavior in subsequent elevated plus maze performance [58].

In combination, the potential of varied antidepressants to upregulate PNN proteolysis and in turn decrease PNN component levels could contribute to reduced PV expression through mechanisms including increased PV-neuron membrane capacitance. Of interest, decreases in the levels of PNN degrading enzymes and increases in protease inhibitors has also been observed in the hypothalamus of obese mice, raising the possibility that excess ECM deposition may occur with different stressors [59].

However, additional non-mutually exclusive mechanisms could also contribute to antidepressant related reductions in PV expression. For example, fluoxetine may disrupt a protein-protein interaction that limits TrkB phosphorylation in PV neurons to in turn reduce PV activity and enhance pyramidal plasticity [60]. Moreover, a separate study also showed that chronic exposure to fluoxetine could upregulate PV surface expression of 5HT2A receptors to in turn reduce PV firing [61].

In summary, these preclinical findings suggest the varied monoamine antidepressant drugs can target PNN components and concomitantly reduce hippocampal PV activity. Importantly, though PNNs predominantly surround PV neurons in the dorsal hippocampus, they can also surround excitatory neurons in hippocampal CA2 select areas of the amygdala [62, 63]. Future studies should therefore examine the effects of varied ADD on additional brain regions as well as PNN dependent effects on plasticity in these regions. Additional preclinical studies could also test more specific PNN modulators, such as the hyaluronan synthesis inhibitor 4-Methylumbelliferone [64], as potential adjuvant treatments for MDD.

## Figure Legends

***Supplemental Figure 1***. Novel tank (fish) and Elevated plus maze (EPM; mice) testing to assess potential correlates of anxiety. The novel tank test is often used with zebrafish to assess anxiety like behavior, with more anxiety typically associated with increased time spent at greater tank depth [65]. As shown in this supplemental figure (upper images) fluoxetine treated fish had a greater latency to reach the bottom of the tank (*p*=0.0007, Student’s *t* test), and also spent more time in the upper portions of the tank during early (first 5 minutes) and later (50-60 minutes) time points. EPM results for mice are shown in the lower image. A previous study showed that venlafaxine treated male mice that were exposed to chronic corticosterone, versus those exposed to saline treated controls exposed to corticosterone, showed less potential anxiety on this test [12]. In this slightly smaller cohort of female non-corticosterone treated mice, however, EPM open arm time did not differ. Viloxazine treated female mice did however show increased open arm time when compared to controls (*p*= 0.0146, Student’s *t* test).

## Notes

### Competing Interest Statement

The authors have declared no competing interest.

### Funding

- National Institutes of Health: AG077002

