## Supplementary figures and images for "Varied monoamine reuptake inhibitors reduce parvalbumin expression; implications for pyramidal cell disinhibition and enhanced neuroplasticity"

### supplemental fig

Supplemental fig. 1

#### Zebrafish

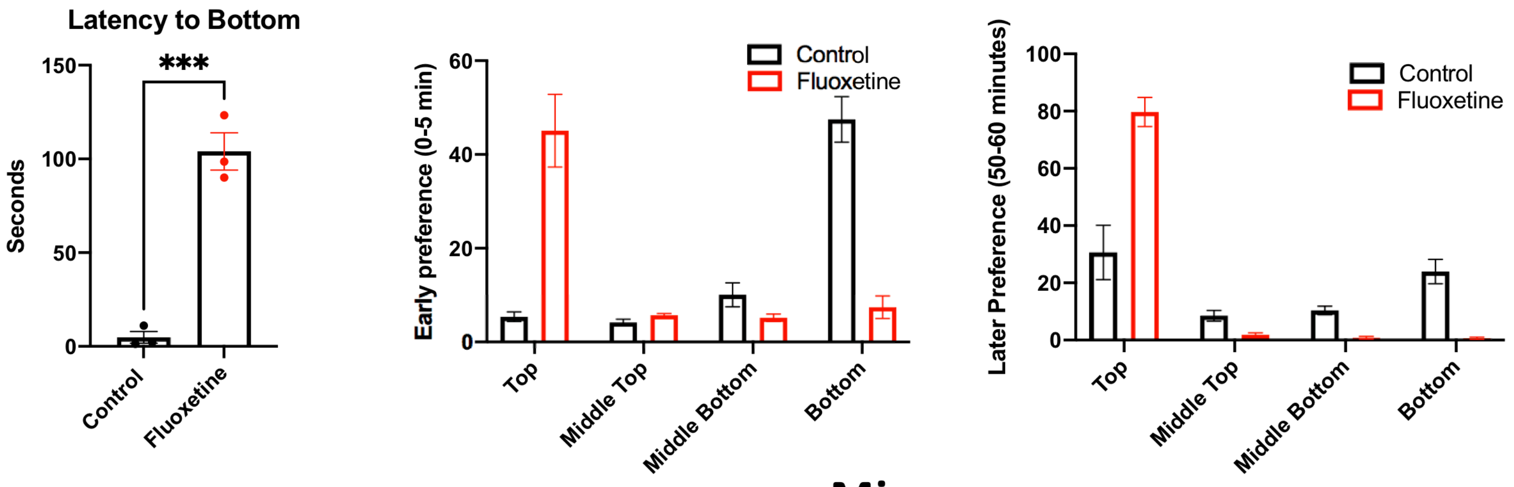


#### Mice

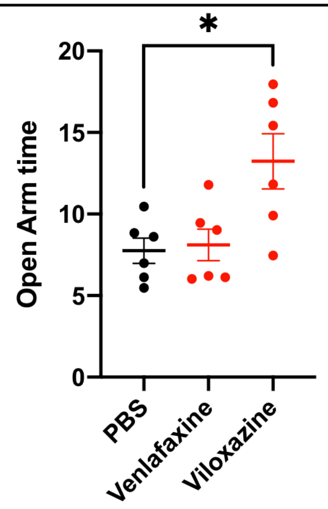
